# OpenEvo: An Open-Source Platform for Automated Evolution and Analysis

**DOI:** 10.64898/2026.07.06.735356

**Authors:** Sebastian S. Cocioba, Pin-Che Huang, John Mallon, Zane Chan, Amanuel W. Geremew, Alex Bisson, Phillip Kyriakakis

## Abstract

Here we introduce OpenEvo, a fully open-source, low-cost turbidostat platform for automated continuous culture and directed evolution experiments. Existing tools are expensive, complex, or lack open-source hardware; OpenEvo addresses this gap with a complete, fully automated evolution platform with detailed, illustrated construction instructions for beginners, open-source software and firmware, priced around $300. An optional PC-based interface offers enhanced functionality, including remote access, programmable evolution cycles, programmable LED stimulation, and a data visualization tool. OpenEvo can cycle through three types of media for positive, negative, and neutral selection conditions, supporting a wide range of experimental designs. We validate the use of OpenEvo by evolving *Haloferax volcanii* to grow from 15% to 12% salt over ∼150 cycles, ∼1,000 hours. Evolved cells grew 55% faster than wild-type at 12% salt. Whole-genome sequencing of adapted cells found SNPs and large deletions. We also demonstrate positive and negative selection using the OpenEvo LEDs to drive optogenetics via a Phytochrome B-based optogenetic tool, with light as the selection stimulus during over 4000 hours of growth. OpenEvo lowers the technical and cost barriers for continuous evolution experiments, serves as a teaching tool, and is designed to grow an open community of users who share modifications.

## Introduction

Continuous culture experiments and directed evolution are powerful approaches for understanding and engineering biological functions. The E. coli Long-Term Evolution Experiment (LTEE), started by Richard Lenski in 1988, exemplifies the utility of long-term growth experiments (Blount et al., 2008, 2012; Tenaillon et al., 2016; Wiser et al., 2013). Similarly, directed evolution typically uses iterative growth and screening, or continuous evolution over many generations, to obtain mutants with improved or novel functions (Wang et al., 2021). A common bottleneck with these tools is the lack of automation and the need for manual passaging of cell cultures (Barrick et al., 2023; Wang et al., 2021). Removing this bottleneck can speed up evolution campaigns or increase throughput by parallelizing experiments. Furthermore, automation tools can enable additional interventions, facilitating new experiments. These interventions include controlled chemical gradients, external stimuli such as light or temperature, or preprogrammed changes in dilution rates and cell densities over time. This open-source automation opens the door to AI-directed evolution experiments, enabling AI to supervise evolution, increase speed, or push the limits of the qualities/functions that can be evolved.

There are a handful of turbidostat/chemostat tools published for lab/research-scale use, but none of them provide guidance on building or assembly methods, and some recent systems can be more complicated than some users need. One tool that enables such automated evolution is the “eVOLVER”, which allows users to grow several cultures in parallel and has open-source software and hardware (Wong et al., 2018; Zhong et al., 2020). However, eVOLVER may be more than some users need in terms of size and throughput (16 parallel cultures), which adds to the effort required for setup and cleanup, and there are no instructions for building one available. A research-scale device called the “Pioreactor” (Gopalakrishnan et al., 2022) is available for purchase, but only the software from the company is open source (the original hardware details are available on GitHub), which limits its flexibility for research labs. Chi.Bio is a feature-rich system that combines the bioreactor with optogenetic stimulation and real-time manipulation of biological systems (Steel et al., 2020). It is available for purchase, sparing users the need to assemble it. However, it is more expensive than OpenEvo and is comparatively complex to build if a user chooses that route for customization or as a teaching tool.

Predating these, other low-cost turbidostats have been published for specific research applications. One device was designed for continuous selection of pooled genetic libraries (McGeachy et al., 2019), and another described a turbidostat that can estimate growth rates in real time (Hoffmann et al., 2017). Guarino *et al*. and Takahashi *et al*. published low-cost turbidostat designs for *in vivo* control experiments and synthetic circuit characterization, respectively (Guarino et al., 2019; Takahashi et al., 2015), but neither includes detailed assembly instructions comparable to OpenEvo’s. Even older devices, while instrumental in the development of these open-source tools, lack modern features and computing capabilities (Bergenholm et al., 2019; Lovitt & Wimpenny, 1981; Matteau et al., 2015; Miller et al., 2013; Stewart & McClean, 2017; Toprak et al., 2013).

While some share build instructions openly, none of these tools provides detailed, illustrated, step-by-step instructions for building, programming, and running the devices. This motivated us to develop OpenEvo and demonstrate that the platform can pass through the challenge of biological validation.

In this manuscript, we describe OpenEvo, a turbidostat device for automated culture growth and laboratory evolution. OpenEvo is a relatively simple device that contains a single growth chamber with hardware and software for monitoring and maintaining growth. All of OpenEvo is open source and modular, making it easy for users to customize for different applications. The cost to build an OpenEvo is ∼$300 for a single machine and $200 for a second one (since some parts come in packs). Here, we present two versions: a standalone version that runs without a PC, and a connected version that runs on a PC with additional monitoring and control features. We demonstrate the magnetic bay’s modularity for clipping on sensors or stimuli, such as LEDs. Up to three media types can be used in a single experiment, such as positive, neutral, and negative selection media. In addition to the device’s features, we developed OpenEvo as a teaching tool. Constructing OpenEvo involves 3D-printing parts, assembling them with computer boards and sensors, installing firmware and software, and cleaning and assembling the fluidics. To make building OpenEvo easier and more approachable, we wrote a detailed and illustrated assembly manual along with separate operating manuals.

Below, we introduce you to OpenEvo, describe its hardware and software features, explain its design, and demonstrate the system’s utility by using it to evolve the halophilic *Haloferax volcanii* to grow under low-salt conditions. We think these tools fill a critical unmet need for studying biology using continuous cultures and can facilitate directed evolution while adding minimal complexity. OpenEvo can be used for teaching in a variety of fields, by hobbyists interested in building or biology, and by research labs that want the flexibility of adopting a completely open-source system. OpenEvo is coded in Python (interface) and Arduino (firmware), so it can be fairly easily modified to add new functions, but no programming experience is necessary to install or run OpenEvo. We hope that the widespread adoption of OpenEvo will lead to an ecosystem in which modifications and forks can be shared openly, fostering innovation and learning.

## Results

### A. Overview of the OpenEvo System

OpenEvo comprises two main components: hardware and software/firmware. The hardware consists of 3D-printed parts, electronics for sensing and control, a fluidics system, and optics for measuring and stimulating cells with light. The software system comprises firmware to operate the hardware and software, enabling the interface to control OpenEvo and support data visualization **(Table1, Figure 1)**.

**Table 1:** Key OpenEvo features.

| Category | Feature / Description |
| --- | --- |
| Core Turbidostat Function | Automated media changes - Triggered when OD exceeds threshold |
|  | Media type tracking - NEUTRAL/POSITIVE/NEGATIVE selection |
| Optical Density Measurement | 940nm IR measurement - TSL2591 light sensor |
|  | 4-point calibration |
| Temperature Control System | Adaptive PID tuning - Aggressive when far, conservative when close |
|  | Heating disabled on overheating - Emergency shutoff prevents damage |
|  | Temperature monitoring - Ambient, media, and heater temps tracked |
| SD Card Data Logging | CSV data format - Standard, easily analyzed format |
|  | UTC timestamps - Real-time clock synchronized from interface |
|  | State restoration - Can resume after power loss |
| LED Cycling for Evolution | 6 LEDs - Cycles through pins 1-6 repeatedly |
|  | PWM brightness control |
| Cycler Program System | Per-step media type - Different media per step (NEUTRAL/POSITIVE/NEGATIVE) |
|  | Per-step LED brightness - Different light intensity per step |
|  | Skip step - Manual advance to next step with media change |
|  | Jump to step - Direct navigation to any step |
| Hardware Control | 4 peristaltic pumps - Adafruit Motor Shield V2 |
|  | Variable pump speeds - Individual PWM control |
|  | Stirring motor - PWM speed control with ramp-up |
|  | Motor safety - Stops on pause/error |
|  | LED driver - TLC5947 for 24-channel PWM |
|  | I2C multiplexer - TCA9548A for multiple sensors |
| Config Commands | 30+ Serial Commands including START, STOP, PAUSE, SET_THRESHOLD, etc. |
| User-Editable Defaults | EMERGENCY_TEMP_LIMIT, incubationSetpointTemp, motorStirringSpeed, etc. |
| Data Integrity | Timestamps, immediate file closing, SD card health monitoring, no pause logging |

**Figure 1:**
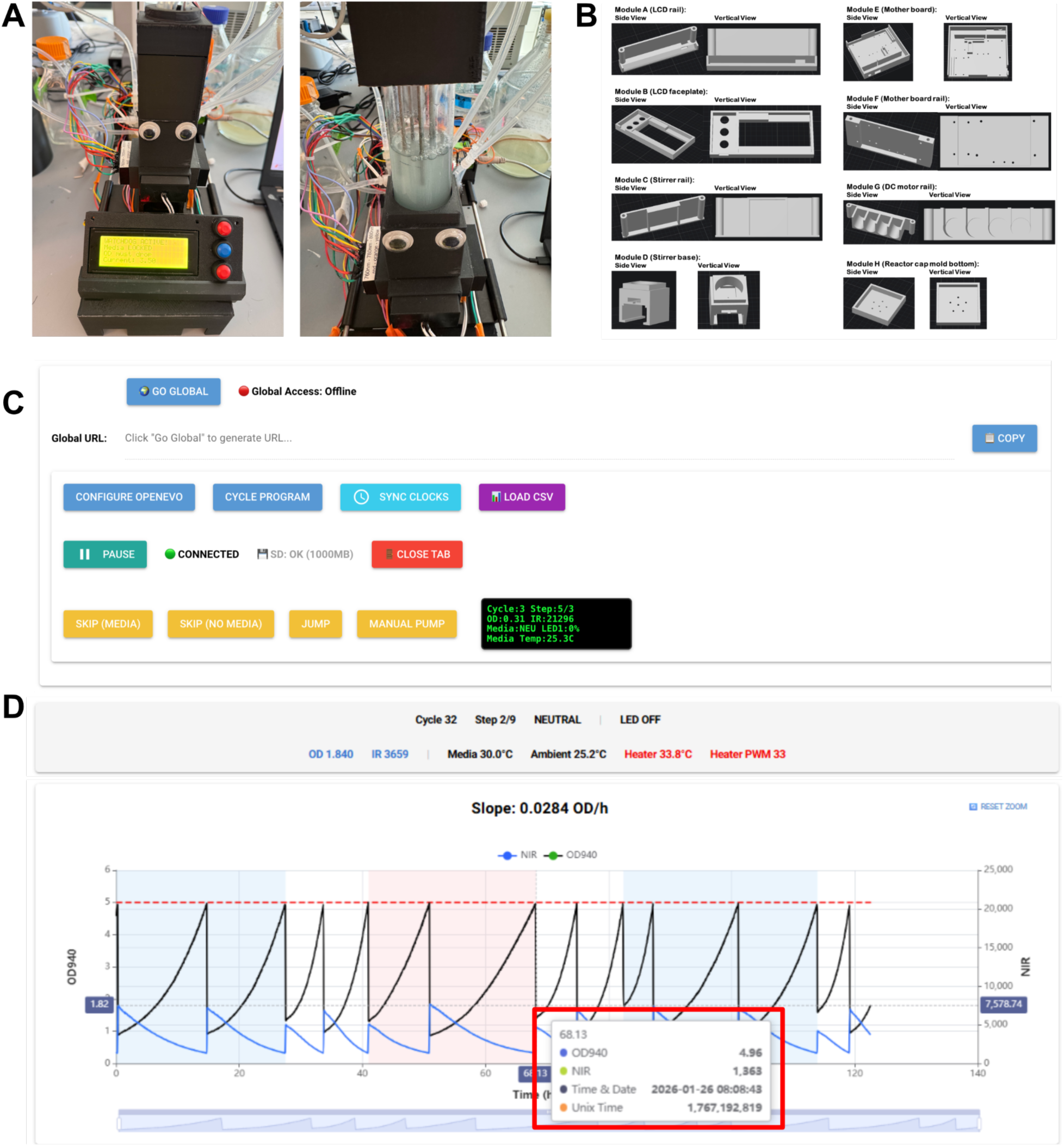
OpenEvo: an open-source and modular robot for automated evolution. **(A)** OpenEvo assembled with optogenetic LEDs running (Left) and open (Right). **(B)** Diagrams of the 3D printed parts used to assemble OpenEvo. **(C)** A screenshot of the web-based user interface control section. **(D)** A screenshot of the status bar, OD/NIR data plotting section of the interface, with a sample of the tooltip (outlined in red) while growing optogenetic Yeast.

At its core, OpenEvo operates as a turbidostat, capable of feeding cells from three different media containers. This design allows for negative and positive selection as well as feeding cells with a non-selective (“Neutral”) medium, though users can use any media (*e.g.*, different selection strengths). OpenEvo uses a 940nm LED to measure the optical density of the cells and a temperature-control system comprising a heater and thermistors to measure the media and heater temperatures. These core turbidostat functions are implemented using an Arduino Mega 2560 microcontroller and its firmware, which controls four peristaltic pumps (three for adding media and one for removing media) and integrates OD940, temperature measurements, and other parameters. OpenEvo was also designed to run without a PC as a standalone device, with all parameters saved to the firmware and an SD card, and designed to support a Graphical User Interface (GUI) **(Figure 1C, 1D)**. The Arduino stores OpenEvo data locally on an SD card to ensure continuous, long-term recording, independent of a computer connection. This data includes periodic timestamps, OD940 readings, temperatures, and more, depending on the firmware version.

The OpenEvo GUI expands capabilities, enabling control of multiple parameters from a PC running Windows or macOS. The GUI features real-time visualization of multiple parameters and the ability to program steps that cycle through various media conditions and/or colored LED illuminations. It additionally provides real-time controls, such as manually pumping media to jump/skip steps and changing configurations. This interface further enables internet-based remote control of OpenEvo from another PC or smartphone. The GUI also supports an OpenEvo module with an LED system for optogenetic stimulation, enabling stimulation with multiple colors using six LEDs on the LED module. LED light dosing can be controlled using Pulse-Width Modulation (PWM) and calibrated through the interface to standard mW/cm2 units.

Within the GUI, there is also a CSV Data viewer for visualizing data from previous experiments **(see Figure 8)**. This viewer provides an easy easily select different experiments in a single CSV file and adjust plotting parameters. It can handle very large files with millions of time points without lagging. We designed the hardware and software to be modular and open-source so that anyone can iterate on new versions or even use parts of it for other applications (*e.g.,* using parts of the interface and hardware to build a simple pump controlled by a GUI). The key features of the OpenEvo system are summarized in **Table 1**.

### B. Hardware and Firmware Workflows

#### System Architecture

To make the platform buildable from scratch for non-experts, we organized all mechanical and electronic parts into modules that users can assemble using our assembly manual. To enable users to build the entire system from scratch, the assembly manual we provide includes complete illustrated assembly instructions, 3D-printed modules, wiring lists, and material sheets.

The hardware features fit into five categories: Sensors & Measurement, Actuators & Controls, the Interface Hardware, Data Storage, and Processing & Communication (**Table 2**). An Arduino controller (Arduino Mega 2560) sits at the core of the system and interfaces with all sensing and actuation modules. The Arduino receives signals from an OD monitor (940nm LED and light sensor) and a thermistor/thermometer. These two monitoring components transmit information back to drive the corresponding actuators, including the heater module, stirring motor, peristaltic pumps, and LED drivers (**Figure 2**). Information from these sensors and states is simultaneously written on an onboard SD card. To enable rapid disassembly and assembly operation, individual modules can be replaced or upgraded without needing to change or disassemble the entire device (**Figure 2**).

**Table 2:**
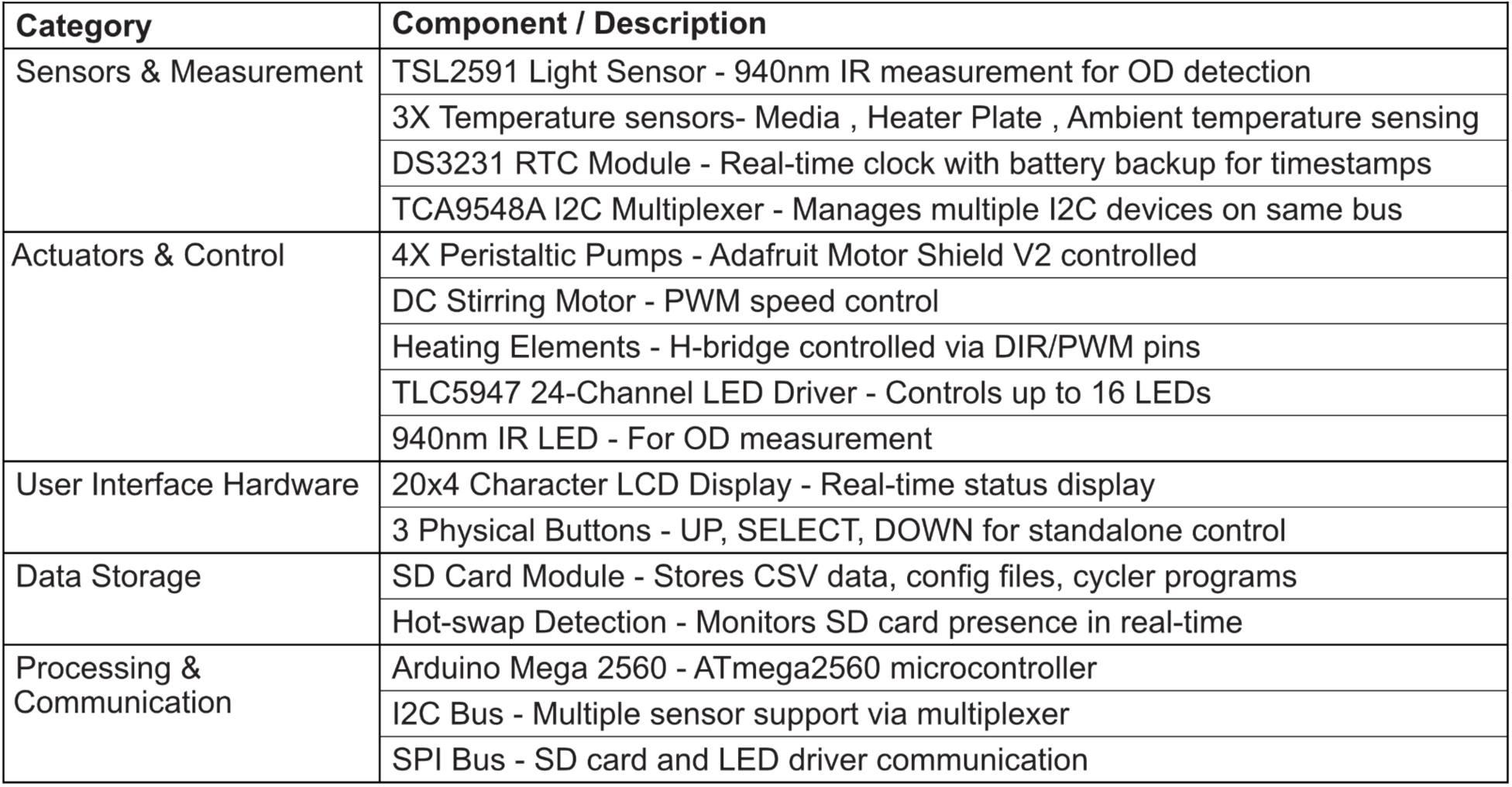
Key Hardware features.

**Figure 2:**
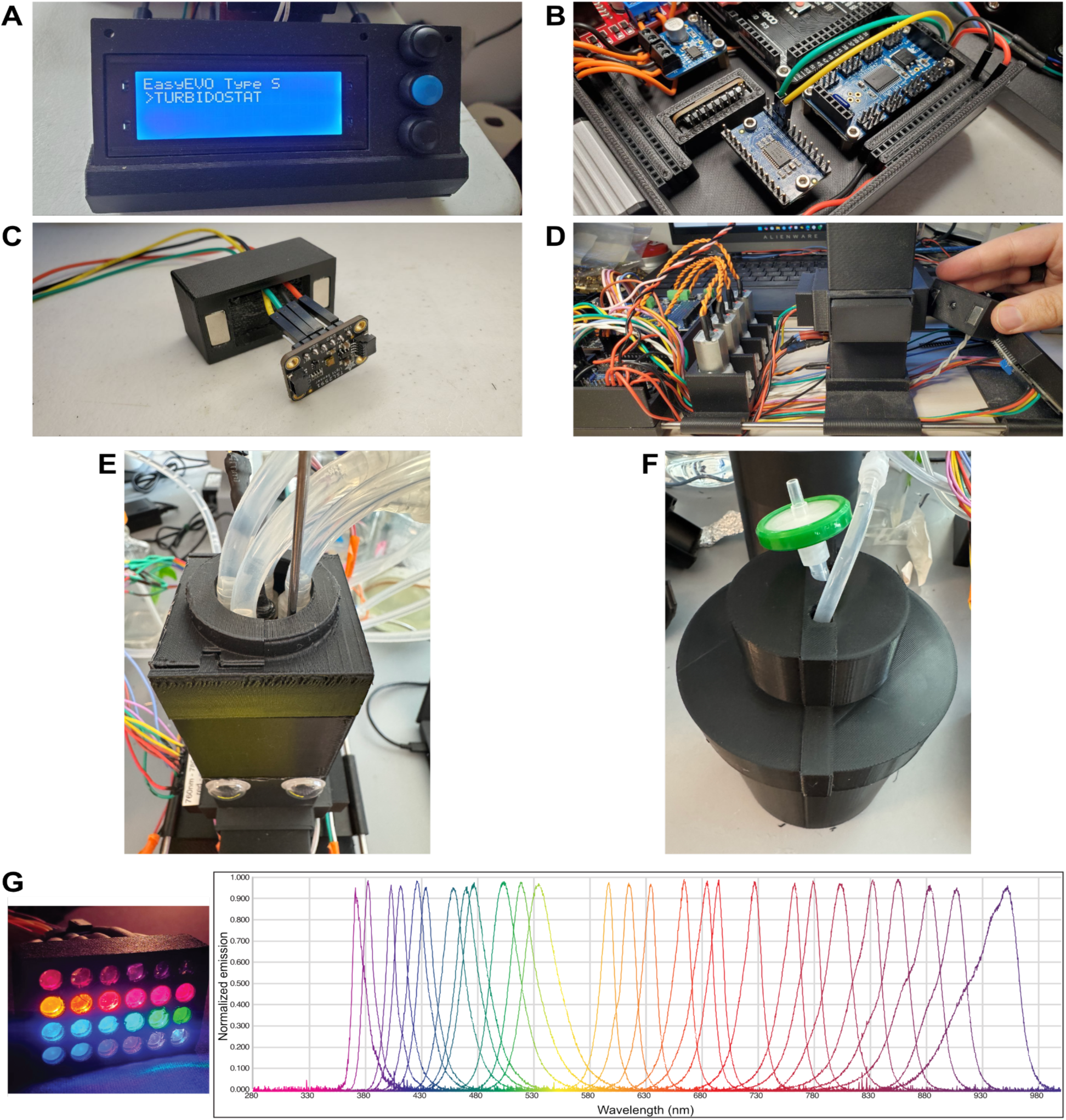
Modular OpenEvo Hardware. **(A)** The OpenEvo LCD interface is mounted using 3D-printed parts. **(B)** The bespoke 3D printed base for holding OpenEvo components. **(C)** The light sensor that measures OD/cell growth is mounted on a modular magnetic mount. **(D)** The 940nm LED for measuring OD is mounted opposite the light sensor, with the growth chamber between them. **(E)** A top to prevent light from entering the growth chamber. Inside is a silicone cap that forms an air-tight seal for the inlets/outlets. **(F)** An accessory for blocking the light from light-sensitive media. **(G)** Inexpensive LEDs and LED emission spectra spanning Violet-NIR.

#### Mechanical Modules

Components of OpenEvo are fabricated using 3D-printed modules designed for easy production and mounting (**Figure 2A-F**). This allows users to easily assemble each component and add custom modules. The main platform has multiple screw holes to secure the electronic components, supporting wiring and mechanical stability. The caps of the culture flasks and media flasks are made of silicone, a highly flexible and heat-resistant material that enables autoclave sterilization and effectively seals the inlet and outlet openings (**Figure 2E**). These are poured into the required custom shapes using 3D-printed molds. The enclosure for the culture flask provides a light-restricted environment that blocks external light from illuminating the cells or the OD monitor (**Figure 2D-E**). Light-blocking containers for media bottles are also available as STL files for 3D printing (**Figure 2F**). A thermistor is placed in a sealed stainless steel needle, providing efficient thermal conduction to the medium. To stir the cells, magnets are attached to a fan that spins a stir bar inside the growth chamber. LED modules with wavelengths spanning the Violet-NIR spectrum can be integrated into the system **(Figure 2G)**.

#### Electronic Connection

The electronic system is organized as shown in **Figure 3**. The automatic decision workflow is then executed via analog input channels on the Arduino controller (Arduino Mega 2560) and its PWM output channels. Users can tune parameters via the physical buttons and LCD display, or update them via the GUI. The SD-card module is connected via the Serial Peripheral Interface (SPI) protocol, enabling high-frequency data logging and “hot-swapping” during experiments, preventing errors when removing the SD card. All wire routing follows the wiring sheet in our manual, and all connections are illustrated, allowing users to assemble or troubleshoot the system without prior electronics experience. Separating analog sensing lines from high-current actuation lines (motors and heater) minimizes interference and ensures stable signal acquisition during long-term evolution experiments.

**Figure 3:**
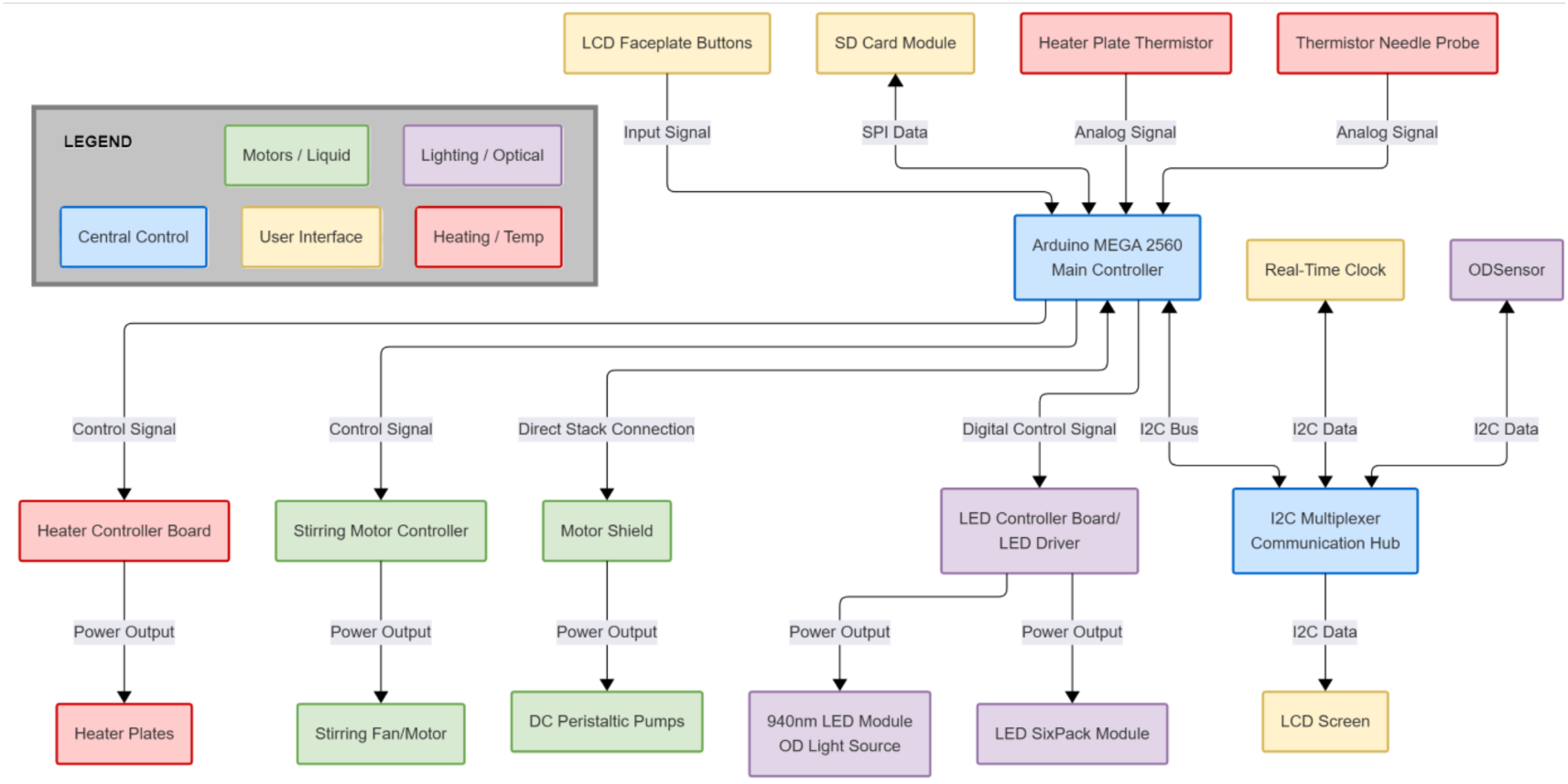
A hardware connection overview of OpenEvo

#### Sterility and Safety Considerations

To maintain sterility, OpenEvo uses a closed-vessel culture system paired with a silicone cap produced using the dedicated cap-mold modules. Every container and tube can be sterilized by autoclaving or bleaching. The silicone cap’s flexibility allows needle insertion for dispensing while maintaining a tight seal. The black cap effectively seals the flask opening to prevent light and contaminants from entering, and a filter on the cap allows the air to move during pumping to prevent pressure buildup. In one instance, we hooked up a pump to continuously bubble air through a filter and sterile water, allowing for constant air exchange without drying or contaminating the culture.

#### Standalone firmware mode

To support OpenEvo in a variety of settings, we made two firmware versions: one that runs all operations on the physical machine without a PC, and another that enables PC control with more advanced features.

In the standalone firmware, all operations are controlled directly on the device using physical buttons and the LCD screen. From the LCD menu, users can run the turbidostat, calibrate the pumps and optics, adjust stirring speed, and set the OD threshold for dilution. In this version, the interface displays only on the LCD module. The standalone version of the firmware allows the physical buttons to be used for parameter adjustments and calibrations. The firmware logs experiment data continuously to the SD card, which users can remove and read on a computer after the run. For users who need only automated turbidostat cycling at fixed settings, without real-time monitoring, remote access, or programmable multi-step cycles, the standalone version offers a lower-complexity option.

#### PC-Controllable and Remotable Firmware

The firmware for running OpenEvo through the interface includes additional features and customizations that make it compatible with the interface. Remote monitoring and control offer numerous benefits, given the time span and the sensitivity of continuous experimental evolution. First, this firmware version includes code to connect to and maintain communication with a PC. Thus, it sends sensor readings periodically and can respond to inputs from the PC interface. The firmware stores default calibrations at the top of the firmware code so they can be easily found and changed by users. Additionally, the interface can be used to change and store non-default calibrations on the SD card. When a new experiment is run, or an old one is continued, the firmware searches the SD card for configuration files before reverting to its defaults, making it easy to continue experiments using the same settings. Because the interface provides a more interactive context, the firmware includes additional functions to enable features such as setting up cycling programs with preprogrammed steps, skipping steps, and performing manual pumping. Importantly, the cycle programs are configured via the interface but stored locally on the SD card, so the firmware can read and run them when disconnected from the PC.

The interface-enabled version of the firmware allows the system to pause and supports “hot swapping” of the SD memory card, which automatically pauses when the card is removed and restarts when it is reinserted. This provides a simple way to download the data and restore the system, keeping it running without creating a new experiment. The data is stored in a format that includes the media type, the cycles and steps, and whether the dilution threshold has been reached. The firmware also includes the ability to control LEDs for optogenetic stimulation and watchdog functions to prevent accidental dilution or overheating. Another major upgrade from the standalone version is OD calibration. The interface version allows for two-, three-, or four-point calibration. Users can pick two points (*e.g.,* upper threshold OD and predicted diluted OD), or more points for a more accurate reading. The OD600 of cells can be measured with a user’s spectrophotometer, and the corresponding OD940 can be obtained from the OpenEvo. The code is annotated extensively to explain how parts like the calibration work.

### C. Buildability / Accessibility

#### Building Manual Summary and Approach to the Building and Teaching Process

Our assembly manual provides detailed step-by-step instructions for building and setting up an OpenEvo from start to finish **(Figure 4-5)**. This includes detailed teaching instructions on tasks such as cutting wires to length, soldering and crimping them, assembling the 3D-printed parts, and wiring everything together. An experienced builder may be able to gloss over some of these parts, but we aimed to make the instructions as complete as possible to make them accessible to non-electronics experts. By having a non-expert follow our instructions to build and set up an OpenEvo, we ensured the instructions were sufficiently complete and also served as a teaching tool for several aspects of the project. Using the assembly manual, an undergraduate student with no prior electronics experience built a functioning OpenEvo with minimal supervision. The team then addressed the obstacles encountered to produce seamless building instructions.

**Figure 4:**
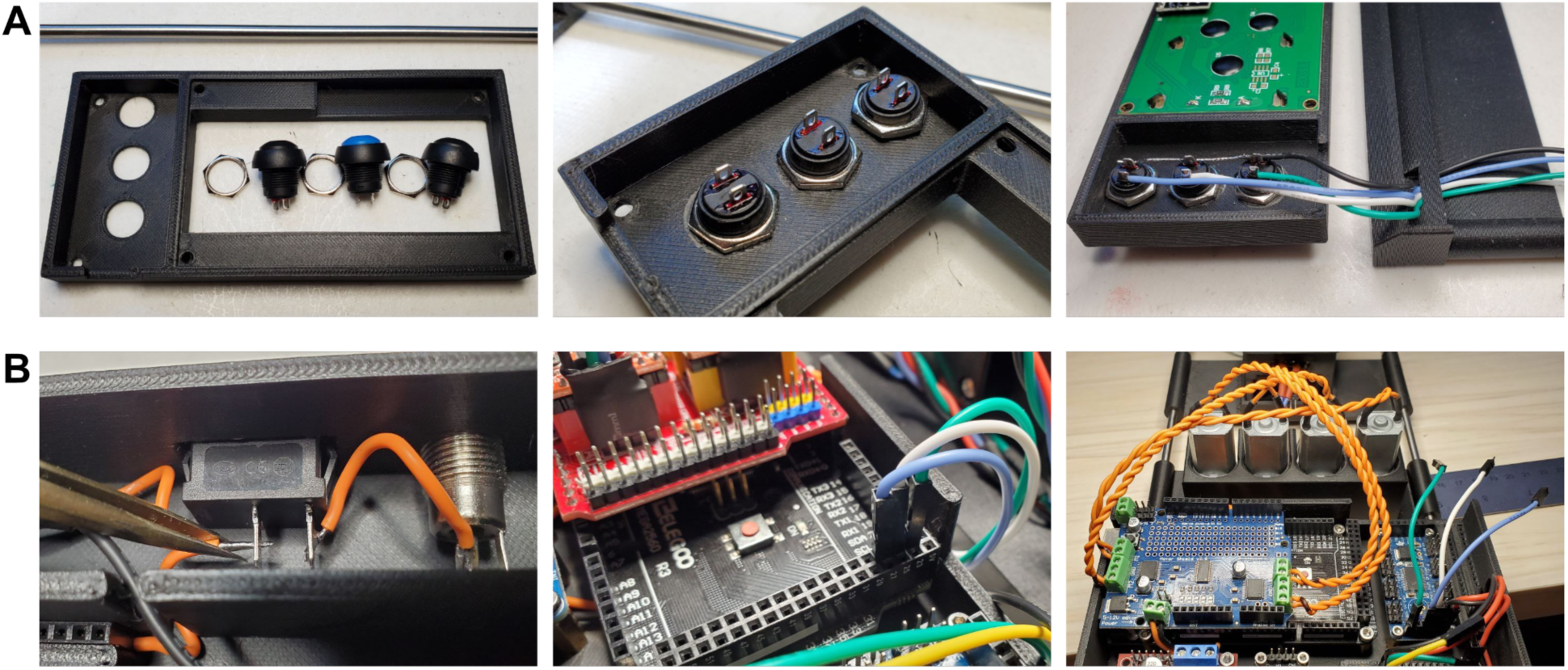
The OpenEvo assembly manual provides step-by-step instructions. **(A)** Example of instructions for assembling 3D-printed parts. **(B)** Pictures from the detailed instructions for wiring OpenEvo using soldering, crimped wire, and screw-in terminal blocks.

**Figure 5:**
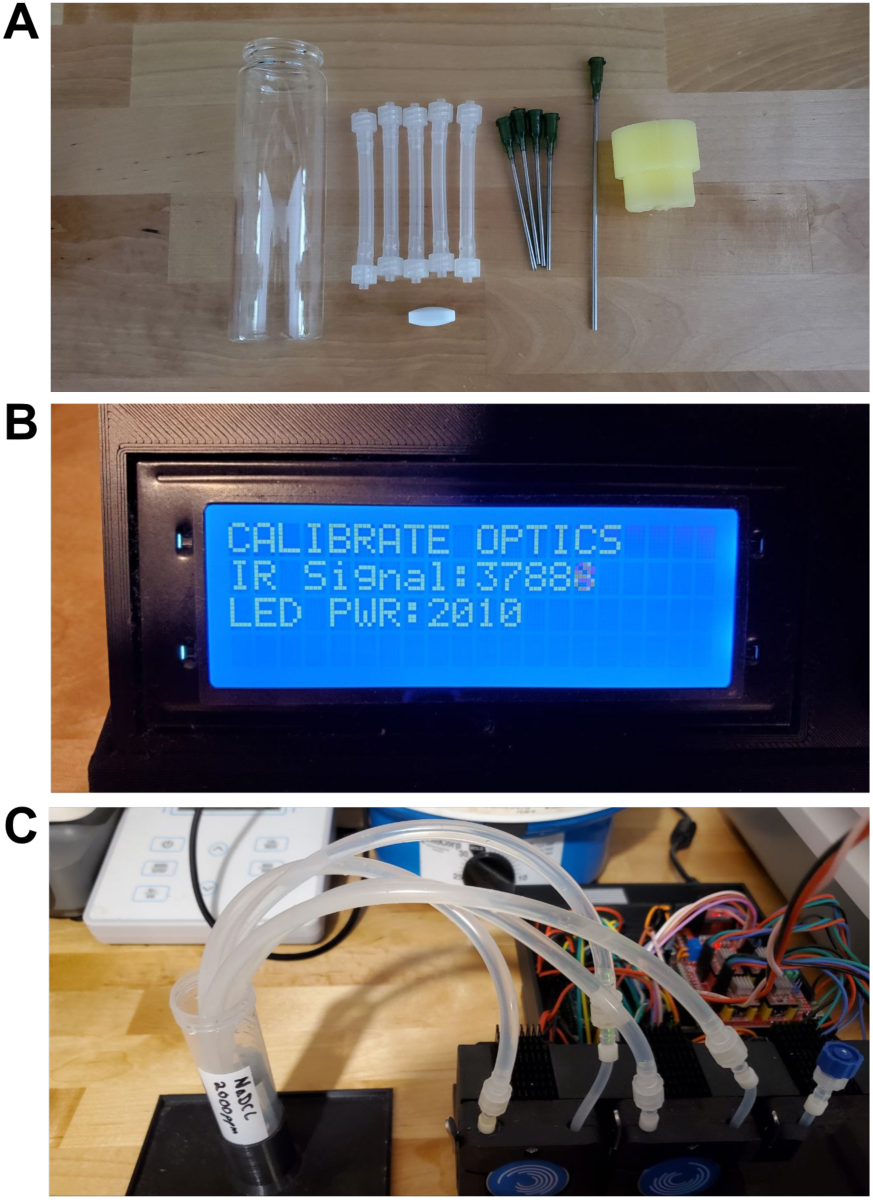
Photographs from the OpenEvo setup manual. **(A)** Parts needed for the reactor vessel prior to sterilization. **(B)** Visual instructions for the calibration setup. **(C)** Instructions for sterilizing OpenEvo.

The resulting assembly instructions include pictures for every step, so builders can check whether their machine matches the instructions. While this makes the instructions longer, it also makes them faster to use and less error-prone. We also compiled the wiring list and material sheet to help users prepare before building a machine. These reduce unnecessary waste and can save considerable time. To run OpenEvo, we created a separate manual for setup and operation once the machine is built. This includes instructions for cleaning and sterilizing the system, calibrating it, and installing the sterile components **(Figure 5)**.

Taken together, we think this project can serve as a tool for self-training to become a “maker” in 3D printing, electronics, and basic software/firmware. While it is impossible to include instructions for every facet of the assembly, such as how to 3D-print the parts using our files, we were as inclusive as possible. We omitted only instructions that are easy to find on YouTube or similar platforms.

### D. Software interface

The user interface software connects a PC to the Arduino via a USB connection **(Figures 1C, 1D, and Figure 6)**. This connection facilitates real-time data output and GUI-based controls of OpenEvo. The interface also lets users connect to the PC over the internet from another PC or a smartphone, with full control of the system. The interface also includes a data visualization feature to open files from previous experiments and explore the data. Key features of the GUI are listed in **Table 3**.

**Figure 6:**
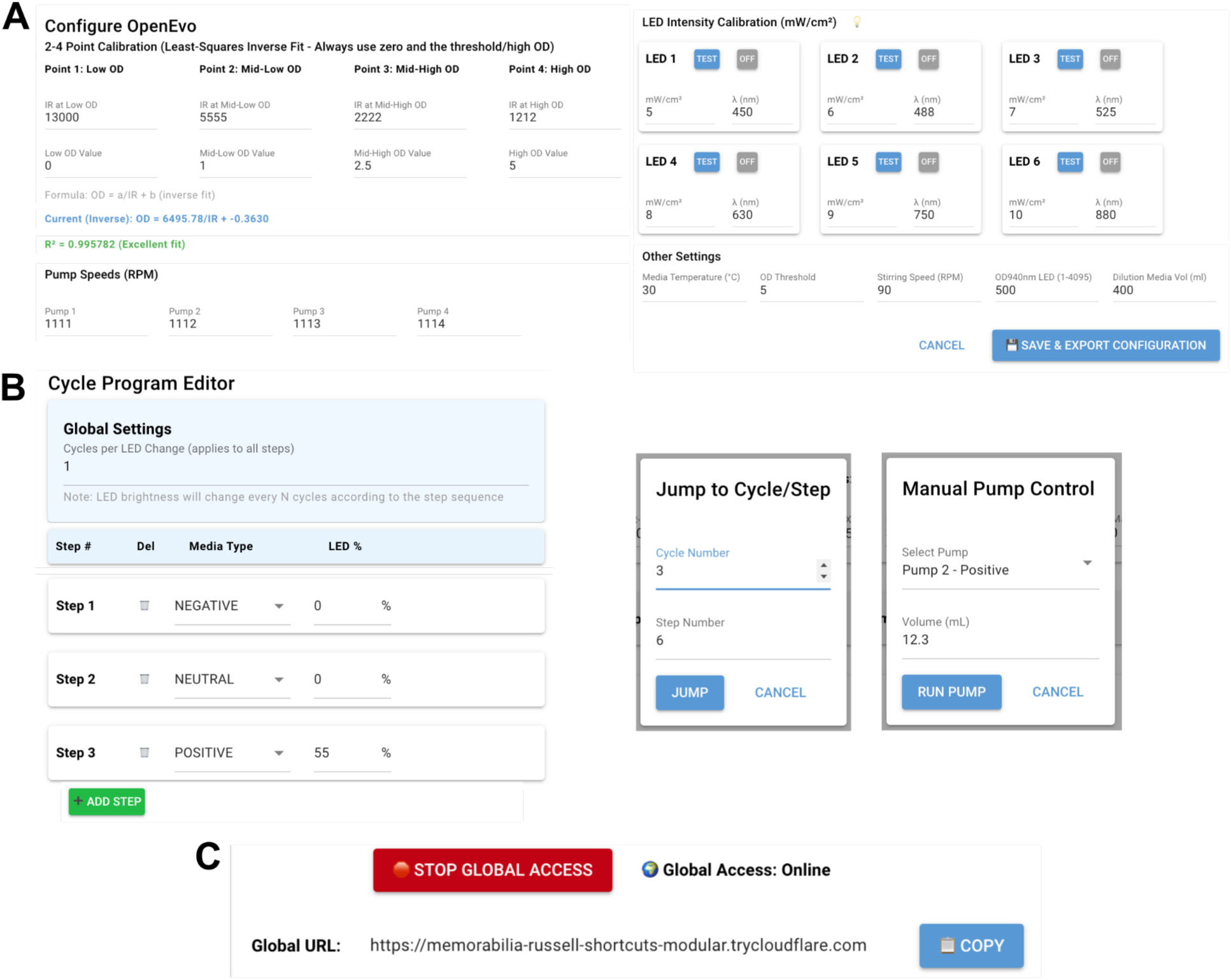
Interface configuration features. **(A)** The configuration window allows the entry of basic setup and calibration parameters. **(B)** The Cycle Program Editor allows you to customize the number of steps per cycle, the media and LED intensity for each step, the number of cycles until the LED changes color, and to add or remove steps. Additional dialogs allow manual skipping, jumping to specific cycles and steps, and manual pumping. **(C)** The interface allows users to create a link to access the OpenEvo interface remotely, which can be used on a computer or smartphone.

**Table 3:** Key Graphic User Interface features.

| *Table 3: Key Graphic User Interface Features* |  |
| --- | --- |
| Category | Feature / Description |
| Connection | Auto-detects Arduino, PING/PONG verification, auto-reconnect |
| Real-Time Display | Live OD/IR/Temperature plots, dual y-axis, zoom/pan, peak detection |
| Data Management | CSV export/logging, SD card access, multi-experiment detection, Unix timestamps |
| Calibration | 2-4-point OD calibration, LED intensity calibration |
| Configuration | Two-click Arduino sync, SD card save and reload |
| Cycler Program | Step editor, media type (NEUTRAL/POSITIVE/NEGATIVE), LED brightness per step |
| Manual Controls | Individual pump testing, precise volume control (0.1mL/pulse) |
| Experiment Control | Start/Stop/Pause/Resume, previous experiment detection, state persistence |
| Status Display | Real-time stats bar (OD, IR, temps, cycle, step, LED intensity), SD card status, LCD mirror |
| CSV Viewer | Opens past experiments, interactive charts, peak analysis, threshold adjustment |
| Access | Local web server (auto-open browser), network accessible |

#### The Interface-OpenEvo Connection

The interface software connects to OpenEvo via the computer’s USB port. The software automatically detects “Arduino-like ports” on a PC or Mac. When you click the connect button, the connection initializes without requiring a port selection. During initialization, the interface detects whether a previous experiment was running on the OpenEvo by reading the SD card. If the interface detects a previous experiment, it will display options to continue the last experiment and to start a new one. This design allows users to pause or disconnect OpenEvo while it is running, and then restart it without changing settings. Once the connection is established and OpenEvo is running, the “CONNECT” button turns into a “PAUSE” button. When paused, no data is recorded, and media changes are not triggered by changes in OD. To close the interface, simply close the browser tab; it doesn’t terminate any connections. The interface can be reopened by entering the same web address. If the PC is disconnected, the connection can simply be restored as described above.

#### Software Requirements and Installation

Version 1 of OpenEvo (Standalone version) requires manual installation of the Arduino IDE software and the required libraries. Version 2 requires installing Python and Arduino, but the dependencies can all be self-installed on a Mac or Windows PC using the installer on the portable zip file. Detailed instructions and files are available on our GitHub repository: https://github.com/Binomica-Labs/OpenEVO/.

#### Configuration and Cycle Program Editor

To configure OpenEvo through the interface, a configuration window lets users enter parameters for measuring and maintaining growth conditions **(Figure 6A)**. Data for the four-point Optical Density (OD940) calibration described below can be entered into this window. The pump speed calibration can also be input in this window. The configuration window further includes options to store LED wavelength and light intensity calibrations **(Figure 6A)**. These calibrations can be performed using the window, with one LED continuously on at 100% brightness, so an external calibrated light meter can record the intensity. The recorded intensity can then be entered into the calibration window, and the calibration will be saved on the SD card for reference. The LEDs are pulsed via PWM to control the light dose. The interface calculates the light dose delivered to the growth chamber, displays it, and stores the calibration data and LED stimulation intensity throughout experiments. These LED data, along with the OD, can be visualized later in our CSV Viewer (see below). Other settings include the media temperature, the OD threshold for triggering cell dilution, the stirring speed, the brightness of the OD940 LED, and the volume of media dispensed during each step. Saving these settings in the interface uploads the parameters to the SD card, allowing OpenEvo to run with those settings even if the PC is disconnected. The settings are also read and saved by the interface when a PC is reconnected to OpenEvo. If users choose to continue the experiment currently being run on OpenEvo, the interface will read the saved parameters and use them to continue. If starting a new experiment, these settings will be the default, but they can be edited prior to beginning a new run. The firmware also has fallback default settings if there is no configuration data on the SD card. The Cycle Program Editor enables OpenEvo users to choose how many cycles trigger a color change of an LED **(Figure 6B)**. Then, the user selects the parameters for each step within the cycles. Steps can be added or removed, and the user can choose a media type, whether the LED is on at that step, and the LED’s brightness. When the save button is clicked, everything is saved as described for the configuration window.

#### Optical Density Calibration Model in Interface-Firmware

The OD module pair uses a 940nm LED and an IR light sensor instead of the typical 600nm light for OD measurement. By using this wavelength, the light used to measure cell density doesn’t overlap with wavelengths typically used in optogenetics experiments or with visible light from ambient laboratory conditions. Depending on the experimental methods and requirements, users may adjust the OD detection range. To this end, we provide two OD conversion models: an Inverse Model (OD = A/IR + B) that fits two to four calibration points, or a simpler two-point Linear Model as an automatic fallback if the inverse model fails.

The Inverse Model is the primary mode for high-precision monitoring as it compensates for non-linear artifacts inherent in measuring the OD at high cell densities. This measurement tracks OD more accurately throughout the growth curve, making it ideal for recording density at different growth points with greater precision. Using two calibration points may not be very accurate at ODs between the two points. However, if the calibration is used to determine the density that triggers dilution as the high point, the user can still ensure the dilution occurs at the correct OD and observe whether the dilution reaches the desired OD. In other words, this can tell users that the OD before and after dilution are happening as expected.

To validate the accuracy of the OpenEvo firmware’s optical monitoring systems, we correlated the IR sensor readings with standard OD600 measurements across a broad dynamic range (OD600 0.2 to 6.0). We initialized three LED intensity settings for IR blank: 32000, 34000, and 37000, using PWM values of 1650, 1850, and 2200 to test under various conditions. The Inverse Model (OD = A/IR + B) shows a high correlation (*R*^2^ ≥ 0.98) between IR and OD600, which verifies our system’s accuracy (**Figure 7**).

**Figure 7:**
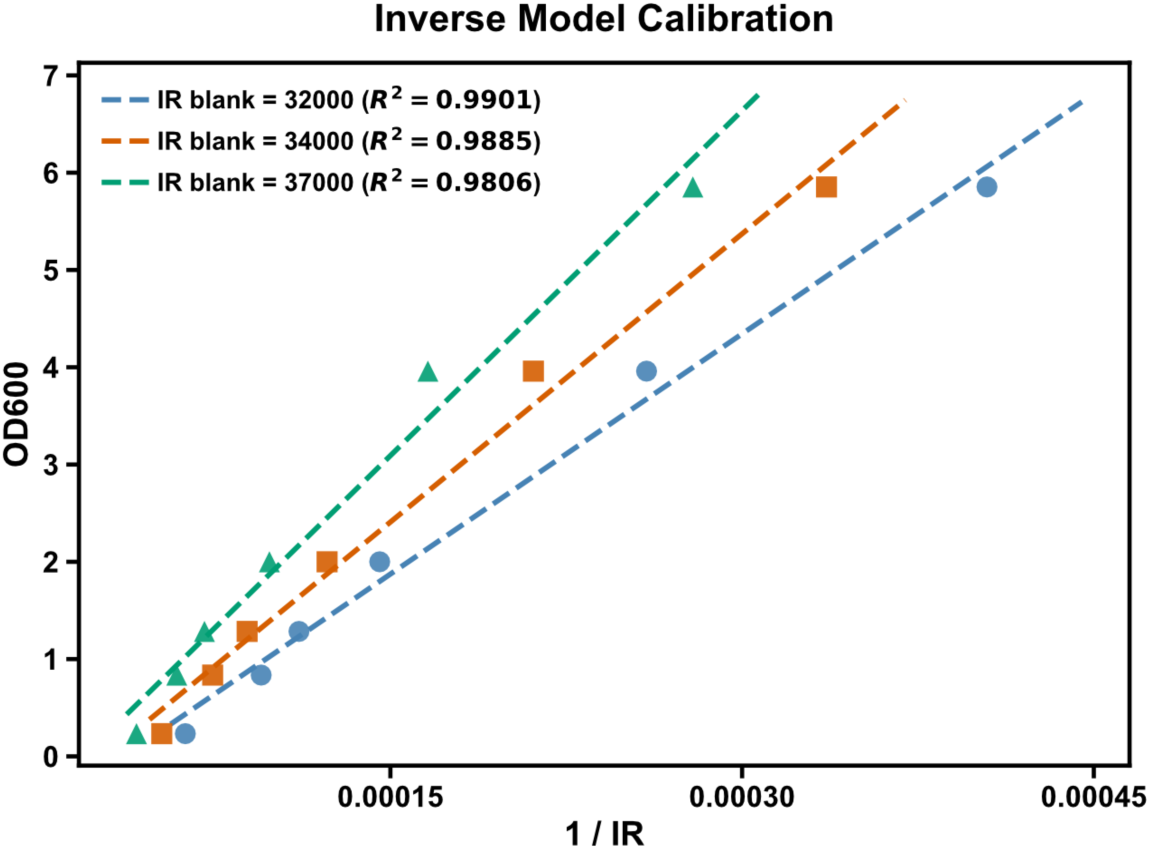
OpenEvo optical density calibration validation. Cells with OD600 values plotted against the r e c i p r o c a l o f t h e I R / 9 4 0 n m L E D r e a d i n g , demonstrating a linear relationship with the inverse calibration algorithm.

#### Real-Time Data Acquisition and Peak-to-Peak Visualization

The Real-Time Data Acquisition interface allows users to visualize several parameters plotted over time **(Figure 6)**. The light detected from the sample using the 940nm NIR LED is plotted alongside the calculated OD600 based on the calibration. The plot’s color shading also indicates the media the cells are growing in. The background of the graph is white, blue, or red for neutral, positive, or negative media, respectively. Below this plot is a graphic that plots the time between peaks. A convenient “Tool Tip” appears when hovering the mouse over the graphs. The tooltip displays the exact time, the OD and NIR values, the date, time, and Unix time. The peak-to-peak distances can be used to assess the growth rate and help quantify the selection pressures being applied. Once the Arduino detects that the OD exceeds the set threshold, it dilutes the cells, sends a message to the interface indicating a peak, and saves the dilution to the SD card. The time differences between these messages are used to plot the peak-to-peak data. Here, the media types are also color-coded with different-colored dots, indicating which media the cells were growing in <u>before</u> they reached the peak. These plots look similar to the ones in the CSV viewer shown in **Figure 8** below.

**Figure 8:**
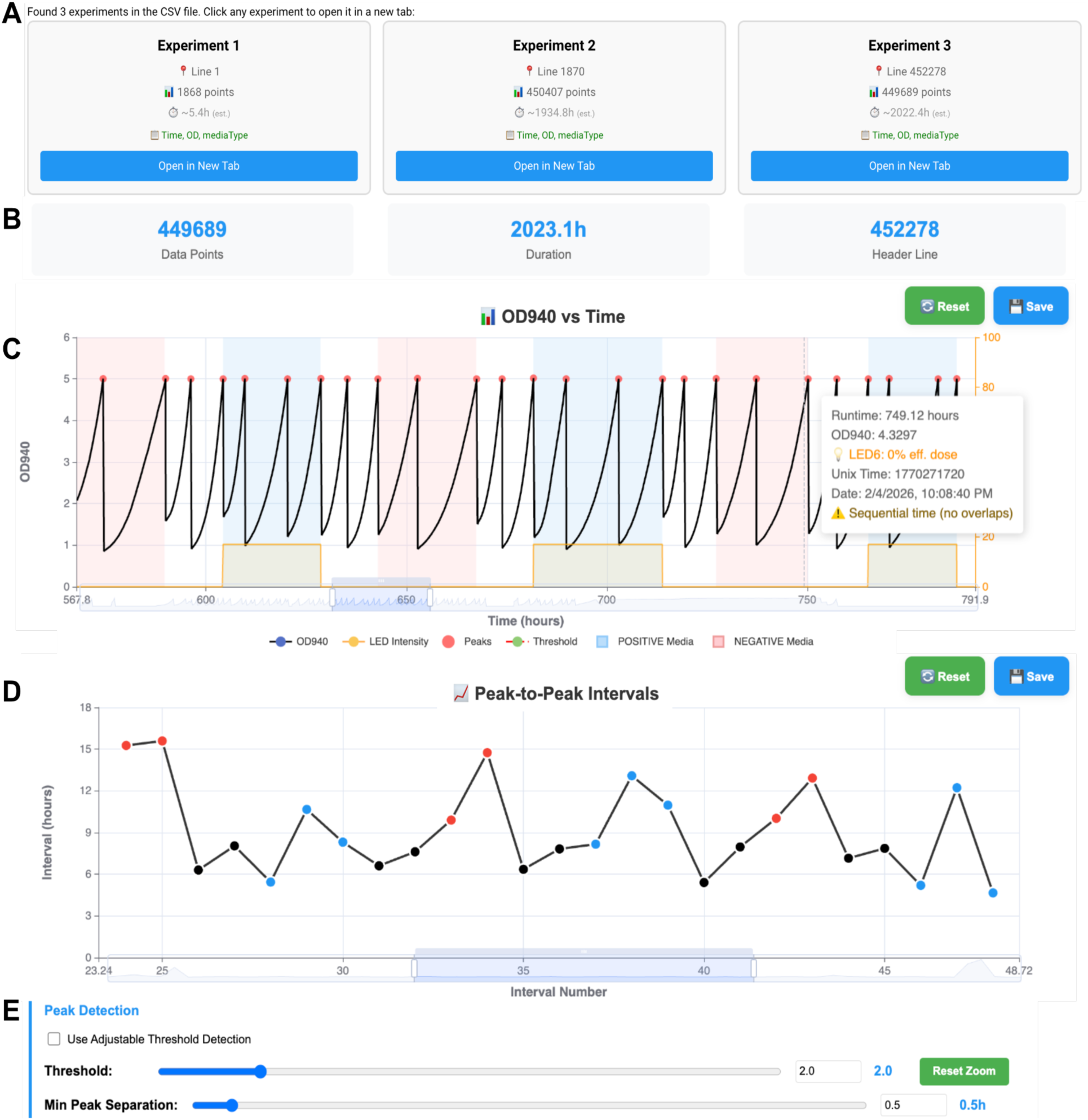
Data viewer for visualizing experiments. **(A)** The CSV Viewer first opens a dialog to select experiments. **(B)** The viewer loads basic statistics and supports axis control, manual control of the peak-detection threshold (optional), and the minimum peak separation (optional). **(C)** The OD viewer shows media types in color. A “tooltip” appears when hovering over the plots to display details about a specific point. LED brightness can be plotted to overlay with the OD data (Orange). The data shown here is from growing the Phytochrome B (PhyB) optogenetic tool in different selection conditions. **(D)** The Peak-to-Peak Interval plot shows the distance between growth peaks, with the points color-coded by media type (Blue = Positive, Red = Negative, Black = Neutral). This data show that applying both positive and negative selection to the PhyB yeast slows culture growth, increasing peak-to-peak distances. **(E)** These plots will update automatically if the Threshold or Min Peak Separation is adjusted.

#### Peak-to-peak Distance Calculations

The peak-to-peak calculation relies on firmware data. During turbidostat mode, the firmware monitor continuously detects the OD value in real time. Once the OD value exceeds the programmed threshold, it activates the dilution pump and simultaneously sends a dilution-event flag and the current media state to the interface. On the interface side, the peak-to-peak interval is calculated using the time differences between consecutive dilution-event flags. This algorithm simplifies peak detection and eliminates noise and misjudgment inherent in local-maxima detection algorithms.. Thus, peak identification corresponds directly to the biological control event executed by the hardware, providing precise observation.

#### Manual Controls and LCD Screen Mirroring

The interface and matching firmware include features for manually controlling several OpenEvo functions. One function allows users to skip a step, including performing the media change that usually occurs between steps. Another function simply changes the program’s state to the next step; no media changes occur. To facilitate moving to any cycle number or step, the jump button lets users select a cycle and step from a drop-down menu **(Figure 6B)**. Additionally, the “MANUAL PUMP” button enables users to select a media type and pump in a specific volume, or to select a volume to empty **(Figure 6B)**. Combined, these functions give users complete manual control of the volumes and media conditions. The interface also has a screen that mirrors the LCD display on the OpenEvo. This helps you monitor what is displayed directly on the OpenEvo hardware or can replace the LCD altogether. This facilitates troubleshooting, especially when working remotely.

#### Go Global: Viewing and Controlling OpenEvo Remotely

Clicking the “GO GLOBAL” button opens a window to create a link to connect to OpenEvo from another internet-connected device **(Figure 1C and Figure 6C)**. The link is copyable via a button, so it can be easily pasted into a web browser or your email. “STOP GLOBAL ACCESS” disconnects the PC connected to OpenEvo through the internet. Additionally, this connection can be used to run the entire system or to view OpenEvo data without installing Python on the remote computer. This connection gives the user complete control of all interface functions described above. This simple protocol allows easy live viewing and remote control of OpenEvo.

#### CSV Viewer

In addition to the real-time view of OD vs. time and peak-to-peak intervals, the interface includes a CSV viewer for viewing previous experiments **(Figure 8)**. The viewer can open very large files quickly and plot them interactively without lag. An option to ignore threshold/dilution events when calculating peaks is available in the CSV viewer. When selecting this option, the threshold OD for peak detection is adjustable, so the peak-to-peak plot updates accordingly. Using the experiment loading window, users can load multiple experiments into different browser tabs.

### F. Biological validation

#### Evolution of Halophilic Archaea to Lower Salt Conditions

As a proof of concept for the standalone version, we employed OpenEvo to evolve the halophilic archaeon *Haloferax volcanii* to grow in reduced-salt conditions. *Hfx. volcanii* typically grows optimally at a total salt concentration of 18% (w/v) (Mullakhanbhai & Larsen, 1975). Exponentially growing wild-type DS2 cells were inoculated into vials containing media with an initial salt concentration of 15% (w/v). To maintain the ionic balance of the medium, all major salts (sodium, potassium, and magnesium) were proportionally diluted. The OD₉₄₀ ceiling and floor parameters were configured to sustain exponential growth, and the culture was incubated at 42°C **(Figure 9A)**. We tracked the slope of each growth cycle to assess deviations in growth rate **(Figures 9B–C)**. At 15% salt, *Hfx. volcanii* maintained a consistent growth rate over ∼12 cycles. The medium was then switched to 12% salt, which led to an immediate and sustained decrease in the growth slope (corresponding to longer cycle times) for approximately 10 cycles, followed by stabilization. After ∼13 additional cycles, the slope gradually increased, suggesting partial adaptation, but it did not recover to the rate observed at 15% salt. Over the next ∼60 cycles, the slope steadily declined again before stabilizing for an additional ∼40 cycles.

**Figure 9:**
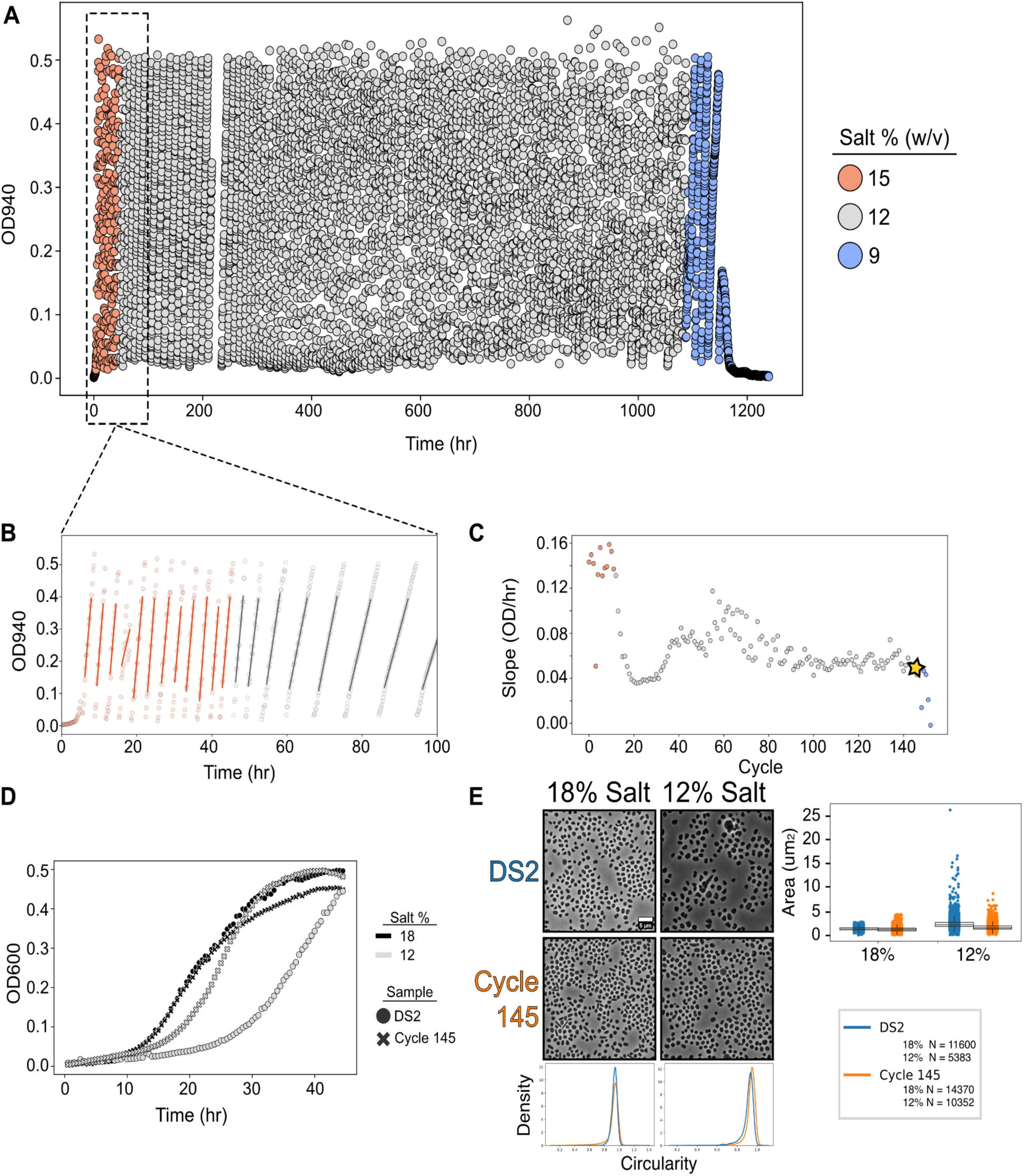
Using OpenEvo to evolve halophilic archaea to grow in lower salt. **(A)** Growth data from the OpenEvo run (graphed every 10th-minute datapoint). **(B)** Zoom in on the first 100 hours to show linear fits that were generated for each cycle. **(C)** Slopes from the linear fits are plotted; cycle 145 is highlighted with a star. **(D)** Growth curves of cycle 145 vs. our wild-type strain (DS2) in both 18% salt (WT conditions) and 12% salt (what cycle 145 was growing in). **(E)** Microscopy pictures and quantification of cells from the different conditions at the end of the growth curves. Scale bar = 5µm.

When the salt concentration was further reduced to 9%, growth progressively slowed and eventually ceased, with the optical density flatlining and the run terminating at cycle 152. Despite the failure to fully restore growth at 12% salt, the culture’s long-term stability over ∼1000 hours suggested some degree of adaptive change. A glycerol stock taken at cycle 145 (evo-145), prior to the 9% salt switch, was revived on agar plates, and individual colonies were compared to wild-type DS2 for growth in both 18% and 12% salt media **(Figure 9D)**.

The evo-145 population grew significantly faster than DS2 at 12% salt, with a doubling time of 4.9 hours versus 7.6 hours (a 55% increase in growth rate). Interestingly, in 18% salt, evo-145 cells grew at the same rate as DS2, with indistinguishable exponential phases and a doubling time of 3.2 hours **(Figure 9D)**, though they reached different carrying capacities. To further evaluate adaptation, we examined single-cell morphologies under both salt conditions. At 12% salt, DS2 cells appeared larger and frequently misshapen, whereas cycle 145 cells were closer to normal size, though slightly rounder. In 18% salt, both strains displayed similar morphologies **(Figure 9E)**.

To map out the genetic variations that could be driving low-salt adaptation of the evo-145 cells at 12% salt, we sequenced a single colony isolated from evo-145 together with the parental DS2 wild-type background from our laboratory collection. Comparison of the evolved isolate to the DS2 control showed three changes in the evolved isolate (Supplementary Table S1). The largest is an 8,838 bp chromosomal deletion (NC_013967.1:2,102,185–2,111,022) that removes eight genes in full, HVO_2243 to HVO_2251. This interval encodes a predicted Mvp-type potassium channel (HVO_2247), a xanthine/uracil permease (HVO_2249), a hypoxanthine/guanine phosphoribosyltransferase (HVO_2250, hpt), and an LctP-family L-lactate permease (HVO_2251). The proximal breakpoint lies within HVO_2242, encoding translation initiation factor aIF-2β, and removes its start codon and first 47 codons; the first in-frame alternative start is a TTG at codon 71, predicting at most a 132-residue N-terminally truncated factor in place of the 202-residue product. The distal breakpoint lies within HVO_2252, an ISH3-like ISH51-family transposase situated at the left boundary of the Halfvol4 provirus (Di Cianni et al., 2025). The remaining two changes are non-synonymous substitutions: Lys117Asn in adenylosuccinate synthetase (HVO_1132, purA), an Mg²⁺- and GTP-dependent enzyme catalyzing the first committed step of AMP biosynthesis, at an allele frequency of 0.74; and Ile340Thr in a DASS-family divalent anion:Na⁺ symporter (HVO_0282) at a frequency of 0.42. Allele frequencies below unity in a clonal isolate are expected in a polyploid organism and indicate that these substitutions are present in a subset of genome copies.

The evolved genotype is therefore compact: loss of an eight-gene operon encoding a predicted potassium channel and two further permeases, truncation of a translation initiation factor, and point mutations in a sodium-coupled symporter and a magnesium-dependent biosynthetic enzyme. All three changes involve either ion handling or the enzymes and machinery that depend on intracellular ion concentrations.

#### Optical Control of Genes for Continuous Directed Evolution of Phytochrome B

Using the interface version of OpenEvo, equipped with LEDs and the cycle-program system, we grew the PhyB-evolving yeast strain under alternating light and dark conditions to apply positive and negative selection. As expected, growth was slower when applying positive and negative selection compared with growth in neutral, non-selection conditions, which served as necessary buffers between the positive and negative selection steps. Our CSV viewer shows these results, displaying the growth curves and peak-to-peak data **(Figure 8)**. Both positive and negative selection initially increase peak-to-peak distance relative to neutral media. Under positive light-driven selection, this slowdown is transient. The intervals shorten during a single selection period, suggesting the population may be adapting to the selection pressure, but sequencing each cycle and characterizing non-evolving clones would be necessary to determine this. (Note that the first step in the positive and negative selections contains ∼10% residual non-selective medium, which can allow more rapid growth than subsequent media changes.)

Because this experiment was conducted during the development of the interface and its firmware/hardware, the dataset contains some gaps. Despite this, OpenEvo maintained growth for over 4,000 hours, including a three-week period when the OpenEvo in California was accessible only remotely from Taiwan. The run continued until the system ran out of media, then resumed after pausing and replacing it. Further analysis will be needed to characterize samples collected throughout the evolution run; nonetheless, this experiment demonstrates OpenEvo’s capacity to run for long periods with minimal hands-on intervention.

## Discussion

We developed OpenEvo after encountering limitations in the accessibility and cost of existing continuous culture systems. As a proof of concept, we ran an evolution campaign of *Hfx. volcanii* through a series of salt reductions over ∼1000 hours, roughly 200 generations, of continuous growth. We found that adapted cells grew faster than the wild-type strain in 12% salt and recovered morphological traits compared to wild-type cells growing at 18% salt. This demonstrated that OpenEvo can run mostly unattended and log growth data in a way that would have been extremely laborious if done by hand. Beyond this experiment, the platform’s modular and open-source design is intended to let others adapt or repurpose the hardware and software for their own workflows.

Our salt-reduction campaign should be read in light of two recent studies that bracket it. Recently, Borst and Soppa established an unevolved *Hfx. volcanii* grows at NaCl concentrations as low as 0.9 M, with roughly 20% of the transcriptome differentially regulated at low salt and small-molecule transport among the classes induced (Borst & Soppa, 2025). While their regime varied NaCl alone, ours diluted all salt components of buffered seawater proportionally, so that our 12% BSW endpoint contains ∼1.6 M NaCl, ∼100 mM Mg2+ and ∼40 mM K+. Our 12% BSW represents, in fact, a potassium concentration roughly 3.5-fold below that of their most extreme low-salt condition. This becomes relevant for an organism that accumulates molar intracellular potassium; proportional dilution imposes a cation-availability problem rather than a sodium problem, and we suspect that the growth arrest we observed at 9% BSW (∼30 mM K+) reflects a potassium rather than a sodium limit.

We envision many extensions and improvements of OpenEvo. What we think would improve OpenEvo the most is a custom Printed Circuit Board that makes it easier to connect the components. While this would make OpenEvo smaller, easier to build, and more organized, builders would have to order a custom Printed Circuit Board, which would limit the system’s accessibility and modularity. Once a user optimizes a modified version and wishes to produce many new versions, they may benefit from a custom Printed Circuit Board that simplifies wiring and other components.

Another limitation in our setup is that it uses a single growth chamber. For some applications, it would be useful to have more than one, particularly for comparing the growth of multiple cultures. One way this could be addressed by using multiple OpenEvos at a time. In the *Hfx. volcanii* experiment, we encountered a lack of controlled airflow through the system. To address this issue, we implemented a separate pump to circulate air through a filter into sterile water (to prevent drying). For our *Hfx. volcanii* experiment, the pump was not very integrated into OpenEvo; it was a separate pump with manual control. Another limitation in OpenEvo is that the interface cannot currently create custom mixes of different media types without manual pumping.

Future expansions or modifications can easily remove these limitations and increase the system’s capabilities. In the future, users could design a custom Printed Circuit Board and make all the files and instructions for ordering and assembling that version available. Adding more growth chambers to an OpenEvo would be a relatively simple modification. At some point, however, the number of inputs and actuators may exceed the current hardware’s capabilities; adding more would require upgrading the hardware and microcontrollers. One workaround is to modify the interface to connect to multiple OpenEvos/Arduinos simultaneously. Similarly, adding more sensors, such as an oxygen sensor with a closed-loop system to pump in air when needed, or a pH sensor and a closed-loop system to buffer or change media, is relatively simple with our platform. Another addition that could be implemented easily without hardware changes is a media pump to deliver air (or specific gases) or to mix media from the three different media inputs. This feature could be implemented by modifying the software to enable cell cultivation under gradually changing media conditions. This powerful addition can be used for a wide array of experiments involving step-up or step-down selection pressure, such as those using salts, nutrients, antibiotics, and more. Another desirable addition would be the ability for OpenEvo to perform controlled mutagenesis, perhaps by illuminating the cells with UV light. This could enable a feedback system that detects when the culture fails to evolve quickly and then induces mutations to help overcome the selection pressure. Similarly, for optogenetic evolution, strains such as the “Dajbog strain” can be implemented with OpenEvo (Gligorovski et al., 2026). We think OpenEvo adopters of OpenEvo will create their own iterations and share new versions with the scientific community for all to use.

## Conclusion

OpenEvo fulfills unmet needs for a cost-effective, modular, fully open-source tool for continuous experimental systems. With accessibility as a primary goal, we developed instructions for building and running OpenEvo so non-experts in electronics or firmware/software can do so. The user interface provides a user-friendly way to interact with OpenEvo, with detailed, highly accessible instructions. We think OpenEvo can serve as a teaching tool for running experiments that are otherwise difficult to perform in a classroom (*e.g.*, passing cells every day) or for teaching how to build devices. It can also be constructed by individuals who want to learn independently and gain a better understanding of the mechanisms by which the machinery and directed evolution operate.

OpenEvo lowers the barrier to entry for automated continuous-culture and directed-evolution experiments. It also allows users to customize the experimental setup because it is entirely open-source. It is also the most affordable modern continuous growth system available, making it more accessible for schools and hobbyists. The illustrated assembly manual likewise facilitates accessibility by eliminating the need for users to interpret sparse text-based instructions.

With OpenEvo, growing continuous cultures for long-term evolution experiments is relatively simple and requires minimal hands-on work, aside from changing media or waste containers. Its internet connectivity also lets users view and control culture growth in real time, regardless of location. Indeed, OpenEvo was showcased in the international 2025 TurBiohacks Bio-hackathon, where students from around the world (*e.g.*, Nepal) controlled an OpenEvo in California.

Using the standalone version of OpenEvo (without the interface), we were able to evolve Halophilic Archaea to lower salt conditions. We found that the evolved cells from the cycle 145 population grew faster than the original strain, DS2, at 12% salt. DS2 cells grew misshapen at 12% salt, whereas cycle 145 cells appeared to have a normal morphology. Genome sequencing revealed many mutations, including some large deletions in the evolved strain. These results raise many new questions about how the cells adapted to the lower-salt environment. We were also able to grow evolving PhyB for thousands of hours, but the analysis resulting from this evolution is beyond the scope of this work. Performing either of these experiments by hand would have been an arduous, less controlled than having OpenEvo perform media changes exactly when cell density reached the prespecified OD.

The current versions of OpenEvo make for versatile research tools, and its fully open-source design is intended to foster a community ecosystem where users can share modifications and accelerate the platform’s utility for the scientific community.

## Materials and Methods

### Building materials

The materials used to construct OpenEvo are listed in the supplemental materials document. The materials list and 3D print/STL files are available on our GitHub (https://github.com/Binomica-Labs/OpenEVO/)

Optional components include a UV-Vis-NIR LED, Black Mica powder (for making black silicone glass vial tops), a pump for air bubbling, a soldering Iron, and helping hands (for holding wires during soldering), all listed in the Bill of Materials available on our GitHub (https://github.com/Binomica-Labs/OpenEVO/).

### Assembly Manual

The step-by-step assembly manual includes all the important information and illustrated diagrams for users to follow, along with notes highlighting common pitfalls and important precautions. Because of the illustrations and large file size, an uncompressed version and a Word Document version (for user editing and note-taking) are hosted on Zenodo rather than GitHub. Zenodo: https://doi.org/10.5281/zenodo.20694531

### Software and Firmware

Software and firmware files are available on GitHub. This includes firmware for the standalone version, firmware for the interface version, and the interface Python file. We also include a portable zip file for self-installation of all required libraries and other dependencies. GitHub: https://github.com/Binomica-Labs/OpenEVO/.

### Strain and Media

The parental strain for the *Haloferax volcanii* experiment was DS2 (Hartman et al., 2010). Cells were grown in the semi-defined, Hv-Cab medium (de Silva et al., 2021), with the only exception being that buffered seawater concentrations were altered by altering the salt-to-water ratios.

The yeast strain used is described in detail in Kyriakakis 2025 (Kyriakakis, 2025). Briefly, yPK-001 had an AH22 background with complete deletions of Leu2, Ura3, Flo1, Met15, and his3delta1 deletion (Scherer & Davis, 1979). In addition, a plasmid (pPK-805) encoding PhyB-Gal4-DBD driven from the pGA promoter (Paulk et al., 2024) was used to express PhyB from the P1 plasmid. PIF3-AD was driven from the HXT7 promoter and PcyA-IRES-Ho1-P2A-FD-P2A-FNR from the TEF1 promoter on plasmid pPK-823. The P1 plasmid was replicated by the BadBoy2 error-prone polymerase (Rix et al., 2024). Sequence files for the plasmids used to make this yeast strain (PhyB = pPK-805, PIF/PcyA = pPK-861) are available on GitHub: https://github.com/Binomica-Labs/OpenEVO/. pPK-805 was transformed into yPK-001 to make yeast strain yPK-063. yPK-063 was then transformed with a PCR product of pPK-861 to make yPK-075. yPK-075 was then transformed with “gr595-pcan1int-iscei-expr-Badboy2sac6p-Natmx” from Rix *et. al.* (Rix et al., 2024) to make yPK-088. yPK-088 was used for the yeast optogenetics experiments. Plasmids and yeast strains are available upon request. Media was composed of YNB (Sunrise Science Cat # 1500-500), Dropout mix (USBiological, Cat # D9540), Histidine (Thermo Scientific, Cat # 411731000), Uracil (Thermo Scientific, Cat # 157300250), with Glutamic acid (Thermo Scientific, Cat # A12919.0B) as a nitrogen source and 2% Dextrose (Criterion, Cat # C5582). For selection, 1mM 3-AT (TCI America, Cat # A0432) and 1mg/ml FOA (Ambeed, Cat # A587031) were added.

## Methods

### LED spectrum measurements

Individual LEDs were measured using a ThorLabs CCS200 spectrometer with a CCSA1 cosine corrector attached to the stock fiber optic cable. LEDs were individually turned on via OpenEvo’s onboard LED driver board and kept at a constant current of 20 mA using a maximum PWM signal from the Arduino microcontroller. The LEDs were measured in direct contact with the cosine corrector’s aperture, and the contact point between each LED and the cosine corrector was temporarily covered with black electrical tape. The spectrum was recorded using the ThorLabs OSA utility software, with a full-spectrum sweep across the CCS200 spectrometer’s range. The resulting data were normalized to the least bright LED and graphed in Excel. A composite of all the waveforms, including the 940nm optical density LED, is shown in **Figure 2G**. The Excel file with the spectral data from each LED is available on GitHub: https://github.com/Binomica-Labs/OpenEVO/.

### OD Transformation Calibration

To monitor OD changes, we use an IR detector to measure the power of 940nm LED light transmitted through the culture to calculate the current OD value. Four OD levels are defined for calibration: Low OD, Mid-low OD, Mid-high OD, and High OD, along with their corresponding IR values from the detector. Cells at different densities were measured using OD600 with a DeNovix DS11 Fx+. Samples above OD600 1.5 were diluted prior to measurement and back-calculated to stay within the accurate range of the Denovix DS11 Fx+. These four OD and IR pairs are entered into the interface, which fits them to an inverse model (OD = A/IR + B) using least squares.

### Using OpenEvo for Optogenetic Control

A vial containing 40 milliliters of media was used as a growth chamber. Three media were used: (1) Neutral Leu-synthetic media, (2) Negative selection media, Leu-media with 1mg/ml 5-Fluoroorotic acid (5-FOA) and 5uM phycocyanobilin (PCB) (Kyriakakis, 2025), and (3) Positive selection media, Leu-Ura-His-media with 5uM phycocyanobilin (PCB) and 1mM 3-amino-1,2,4-triazole (3-AT) (Kyriakakis, 2025). Cells were grown with the 650nm LED to induce URA3 and His3 expression, permitting light-responsive cells to grow under the positive selection. Cells grown in Neutral or Negative selection media were kept in the dark. The OD940 threshold for dilution was set to 5.0.

### Adaptive Growth of Hfx. volcanii cultures in OpenEvo

A flask containing 50 milliliters of HvCab medium was inoculated with a single DS2 colony and incubated overnight. Cells were then back-diluted to an OD600 of 0.05. When the culture reached an OD600 of 0.5, 5 mL was used to inoculate an OpenEvo growth vial containing HvCab media with 15% (w/v) buffered seawater salt. The media used in the OpenEvo device were supplemented with 10 micrograms of carbenicillin per milliliter to prevent bacterial contamination throughout the experiment. An OpenEvo run was started with the temperature set to 42°C, an OD940 ceiling of 0.5, and a floor of 0.003. An aquarium bubbler was also installed, with an indirect connection to the OpenEvo flask via a Schott bottle of autoclaved water heated to 37°C to maintain proper humidity and oxygen levels (de Silva et al., 2021). Growth was monitored for a set period, and then the media bottles at the inlet were swapped to lower the salt concentration to the next level. This process was repeated until the experiment was terminated.

### Comparing cycle evo-145 growth to DS2

Agar streaks of DS2 or cycle evo-145 were made on medium containing 18% or 12% (w/v) Buffered salt water (BSW) (de Silva et al., 2021). Colonies were picked and grown overnight, then back-diluted to an OD600 of 0.05. Cells were then incubated to an OD600 between 0.3 and 0.5 before inoculating a twelve-well plate to an OD600 of 0.05 in 1.5 milliliters of media, and the plate was grown in an EPOCH plate reader (BioTek) at 42°C until the carrying capacity was reached. Wells containing HvCab media at 18% salt and 12% salt were used for blank subtraction. Exponential fits were then made to the region of the curve preceding the inflection point. Doubling times were calculated by dividing the natural log of 2 (ln(2)) by the rate extracted from fitted curves (µ). Graphs are averages of 3 independent experiments, each with its own averaging of two technical replicates.

### Single-cell morphology of cycle evo-145 and DS2 parental strain

Ten microliters of cells from the end of the twelve-well plate growth experiments were placed on a number 1.5 coverslip, immobilized with a 0.5% (w/v) agarose pad at the respective salt concentration, and imaged at 42°C using a Nikon TI-2 inverted microscope within an Okolab H201 enclosure. Phase-contrast images were acquired with a Hamamatsu ORCA Flash 4.0 v3 sCMOS camera (6.5 microns/pixel), a CFI PlanApo Lambda 100x DM Ph3 Objective. Acquisition parameters were 200 milliseconds at 100% power. Representative crops of the data were then selected and placed in the figure.

### Illumina whole-genome sequencing of the evo-145 strain

Agar streaks of the evo-145 population were made on Hv-Cab adapted medium containing 12% (w/v) BSW salt. A single colony was used to inoculate an Hv-Cab culture containing 12% BSW, and genomic DNA was isolated at stationary phase using the Zymo DNA Miniprep Kit. Two DS2-background control strains from the laboratory collection were prepared in parallel. Libraries were prepared by SeqCenter (Pittsburgh, PA) using the tagmentation-based Illumina DNA Prep kit with custom IDT 10 bp unique dual indices and a target insert size of 280 bp, without additional fragmentation or size selection. Sequencing was performed on an Illumina NovaSeq X Plus, generating 2 × 151 bp paired-end reads. Demultiplexing, quality control, and adapter trimming were performed with bcl-convert v4.2.4. Mean read depth was 141× for the evolved isolate and 70–102× for the controls. Coverage was then extracted and plotted using a custom Python script. Raw sequence reads are available under accession number XXXXXX and on Zenodo repository for download: https://doi.org/10.5281/zenodo.20694531.

### Evo-145 variant calling and filtering

Reads were analyzed with breseq v0.40.2 against the H. volcanii DS2 reference genome (RefSeq GCF_000025685.1; accessions NC_013967.1, NC_013968.1, NC_013965.1, NC_013964.1, NC_013966.1), using consensus mode for the controls and polymorphism mode for the evolved isolate, since Hfx. volcanii is polyploid and a clonal isolate may carry a variant in a subset of genome copies. Control call sets were subtracted from the evolved call set with gdtools; variants shared with either control were treated as reference-genome discrepancies or as pre-existing differences in the parental laboratory lineage rather than as products of the evolution experiment. For copy-number and deletion analysis, reads were aligned with BWA-MEM v0.7.17 and per-base depth computed with SAMtools v1.19 at a mapping-quality threshold of MAPQ ≥ 30. Deletion endpoints were defined as the outermost positions with zero coverage at MAPQ ≥ 30. Strain identity of all sequenced isolates was confirmed by read coverage over pHV2 and pyrE2.

## Supporting information

Supplementary_Table_S1_mutations

## Acknowledgements

The authors would also like to acknowledge Vincent J. Hu and Chang C. Liu for many helpful discussions and for supplying the reagents to express the PhyB-PIF3 optogenetic system in yeast containing the P1 plasmid. Finally, the authors would like to acknowledge the funding awarded to P.K.: NIGMS R21 GM143806-01. This work was supported by grants awarded to A.B.: NIH-NIGMS R35 MIRA (7R35GM156992) and the Brandeis National Science Foundation (NSF) Materials Research Science and Engineering Center (MRSEC) Bioinspired Soft Materials (NSF-DMR 2011846). A.B. is the recipient of an NSF grant (NSF-MBC 2222076) and is a Pew Scholar in the Biomedical Sciences, supported by The Pew Charitable Trusts.

## Abbreviations

BSW: Buffered salt water
CSV: Comma-separated values
DC: Direct current
GUI: Graphical user interface
I2C: Inter-integrated circuit
IDE: Integrated development environment
IR: Infrared
LCD: Liquid-crystal display
LTEE: Long-Term Evolution Experiment
NIR: Near-infrared
OD: Optical density
PCB: phycocyanobilin
PhyB: Phytochrome B
PID: Proportional-integral-derivative
PWM: Pulse-width modulation
RTC: Real-time clock
sCMOS: Scientific complementary metal-oxide-semiconductor
SPI: Serial peripheral interface
STL: Stereolithography
UTC: Coordinated universal time
UV: Ultraviolet
WT: Wild type

