## Supplementary_Table_S1_mutations for "OpenEvo: An Open-Source Platform for Automated Evolution and Analysis"

**Supplementary Table S1.** Mutations specific to the evo-145 evolved isolate of *Haloferax volcanii*.

| **Mutation Class** | **Replicon** | **Coordinates (bp)** | **Locus and Annotation** |
| --- | --- | --- | --- |
| Deletion | Main Chromosome | 2,102,185 – 2,111,022 | HVO_2243–HVO_2251 (8 genes) deleted in full, including HVO_2247 (Mvp-type potassium channel superfamily protein), HVO_2249 (xanthine/uracil permease family), HVO_2250 (hpt, hypoxanthine/guanine phosphoribosyltransferase) and HVO_2251 (LctP-family L-lactate permease). Breakpoints fall within HVO_2242 (aIF-2β) and HVO_2252 (ISH3-like ISH51-family transposase). |
| SNP | Main Chromosome | 1,032,839  C→G | HVO_1132 (purA), adenylosuccinate synthetase (EC 6.3.4.4); Mg²⁺- and GTP-dependent. First committed step of AMP biosynthesis. |
| SNP | Main Chromosome | 252,878  A→G | HVO_0282, DASS-family (divalent anion:Na⁺ symporter, SLC13/TC 2.A.47) transport protein, 582 aa. |
